# Shared Data Science Infrastructure for Genomics Data

**DOI:** 10.1101/307777

**Authors:** Hamid Bagher, Usha Muppiral, Andrew J Severin, Hridesh Rajan

## Abstract

**Background:** Creating a computational infrastructure to analyze the wealth of information contained in data repositories that scales well is difficult due to significant barriers in organizing, extracting and analyzing relevant data. Shared Data Science Infrastructures like Boa can be used to more efficiently process and parse data contained in large data repositories. The main features of Boa are inspired from existing languages for data intensive computing and can easily integrate data from biological data repositories.

**Results:** Here, we present an implementation of Boa for Genomic research (BoaG) on a relatively small data repository: RefSeq’s 97,716 annotation (GFF) and assembly (FASTA) files and metadata. We used BoaG to query the entire RefSeq dataset and gain insight into the RefSeq genome assemblies and gene model annotations and show that assembly quality using the same assembler varies depending on species.

**Conclusions:** In order to keep pace with our ability to produce biological data, innovative methods are required. The Shared Data Science Infrastructure, BoaG, can provide greater access to researchers to efficiently explore data in ways previously not possible for anyone but the most well funded research groups. We demonstrate the efficiency of BoaG to explore the RefSeq database of genome assemblies and annotations to identify interesting features of gene annotation as a proof of concept for much larger datasets.

## Background

The amount of data that can be generated by a single sequencing machine in a single day continues to grow exponentially every year [1]. With this data, researchers are able to ask more complex questions in biology. Unfortunately, these data sets have significant barriers for researchers in organizing, extracting and analyzing relevant data to test hypotheses in addition to the barriers for implementing scalable computing infrastructure. The time required to perform data wrangling tasks is a well-known problem in bioinformatics [2] that increases with amount of data. As we scale up the number of files and metadata used in an analysis, a more robust system for reading, writing and storing data will be needed.

This can be achieved by borrowing methods and approaches from computer science. Boa is a language and infrastructure that abstracts away details of parallelization and storage management by providing a domain specific language and simple syntax [3]. The main features of Boa are inspired by existing languages for data-intensive computing. These features include robust input/output, querying of data using types/attributes and efficient processing of data using functions and aggregators. Boa can be implemented as a Shared Data Science Infrastructure (SDSI). Running on a Hadoop cluster [4], it manages the distributed parallelization and collection of data and analyses. Boa can process and query terabytes of raw data. It also has been shown to substantially reduce programming efforts, thus lowering the barrier to entry to analyze very large data sets and drastically improves scalability and reproducibility [4]. Raw data files are described to Boa with attribute types so that all the information contained in the raw data file can be parsed and stored in a binary database. Once complete, the reading, writing, storing and querying the data from these files is straightforward and efficient as it creates a dataset that is uniform regardless of the input file standard (GFF, GFF3, etc) The size of the data in binary format is also smaller.

Genomics-specific languages are also common in high-throughput sequencing analysis such as S3QL, which aims to provide biological discovery by harnessing Linked Data [5]. In addition, there are libraries like BioJava [6], Bioperl [7], and Biopython [8] that provide tools to process biological data.

There are also several tools in the field of high-throughput sequencing analysis that use the Hadoop and MapReduce programming model. Heavy computation applications like BLAST, GSEA and GRAMMAR have been implemented in Hadoop [9]. SARVAVID [10] has implemented five well-known applications for running on Haddop: BLAST, MUMmer, E-MEM, SPAdes, and SGA. BLAST [11] was also rewritten for Hadoop by Leo *et.al.* [12]. In addition to these programs, there are other efforts based on Hadoop to address RNA-Seq and sequence alignment [13, 14, 15].

Unfortunately, there currently does not exist a tool that combines the ability to query databases, with the advantage of a domain specific language and the scalability of Hadoop into a Shared Data Science Infrastructure for large biology datasets. Boa on the other hand is such a tool but is currently only implemented for mining very large software repositories like GitHub and Sourceforge.

There are several very large data repositories in biology that could take advantage of a biology specific implementation of Boa: The National Center for Biotechnology Information (NCBI), The Cancer Genome Atlas (TCGA), The Encylopedia of DNA Elements (ENCODE). NCBI hosts 45 literature/molecular biology databases and is the most popular resource for obtaining raw data for analysis. NCBI and other web resources like Ensembl are data warehouse for storing and querying raw data, sequences, and genes. TCGA contains data that characterizes changes in 33 types of cancer. This repository contains 2.5 petabytes of data and metadata with matched tumor and normal tissues from more than 11,000 patients. The repository is comprised of eight different data types: Whole exome sequence, mRNA sequence, microRNA sequence, DNA copy number profile, DNA methylation profile, whole genome sequencing and reverse-phase protein array expression profile data. ENCODE is a repository with a goal to identify all the functional elements contained in human, mouse, fly and worm. This repository contains more than 600 terabytes (personal communication with @EncodeDCC and @mike_schatz) of data with more than 40 different data types with the most abundant data types being ChIP-Seq, DNase-Seq and RNA-Seq. While it is common to download and analyze small subsets of data (tens of Terabytes for example) from these repositories, analyses on the larger subsets or the entire repository is currently computationally and logistically prohibitive for all but the most well funded and staffed research groups.

Here we discuss an initial implementation of Boa for Genomics (BoaG) that mines the data and metadata contained in the 97,716 genome annotation files (GFF) from the RefSeq database which is subset of the data contained in the NCBI repository. We show how summary statistics of clades of genome assemblies and annotations can provide insights genome/annotation quality, variability in genome quality based on assembler program and unexplored biology of exon frequency in gene models.

## Methods

### Choice of Biological repository for prototype implementation

For our initial implementation of BoaG, we chose to showcase Boa’s database querying ability on RefSeq data. RefSeq is a comprehensive, integrated, non-redundant, well-annotated set of sequences that spans the tree of life containing species of plants, animals, fungi, archaea and bacteria. RefSeq also has a decent amount of metadata about genome assemblies and their annotations for which as far as we know has not been explored as a whole. Once inside our BoaG infrastructure it is relatively straightforward to ask questions that would be challenging to answer directly from the online respository.

- What are the top ten Eukaryotic genomes with the highest and lowest average exons/gene?
- What are the top five most used assembly programs for Insects? How well do they assemble genomes?
- What are the top five most used assembly programs for Bacteria? Is it different than Insects?
- What is the smallest and largest genome in RefSeq?

### Design goals and implementation

In this section, we will provide an overview of the implementation of Boa for the Biology domain which we will refer to as BoaG. This includes its domain specific language design and Hadoop based infrastructure implementation. The overall workflow for BoaG requires a program written in BoaG that is submitted to the BoaG infrastructure, as seen in Figure 1(a). The infrastructure takes the submitted program and compiles it by the BoaG compiler and executes the program on a distributed Hadoop cluster using a BoaG formatted database of the raw data. BoaG has aggregators which are functions that run on the entire database or a large subset of the database and therefore takes advantage of the BoaG’s database designed for both the data and the compute to be distributed across a Hadoop cluster.

**Figure 1:**
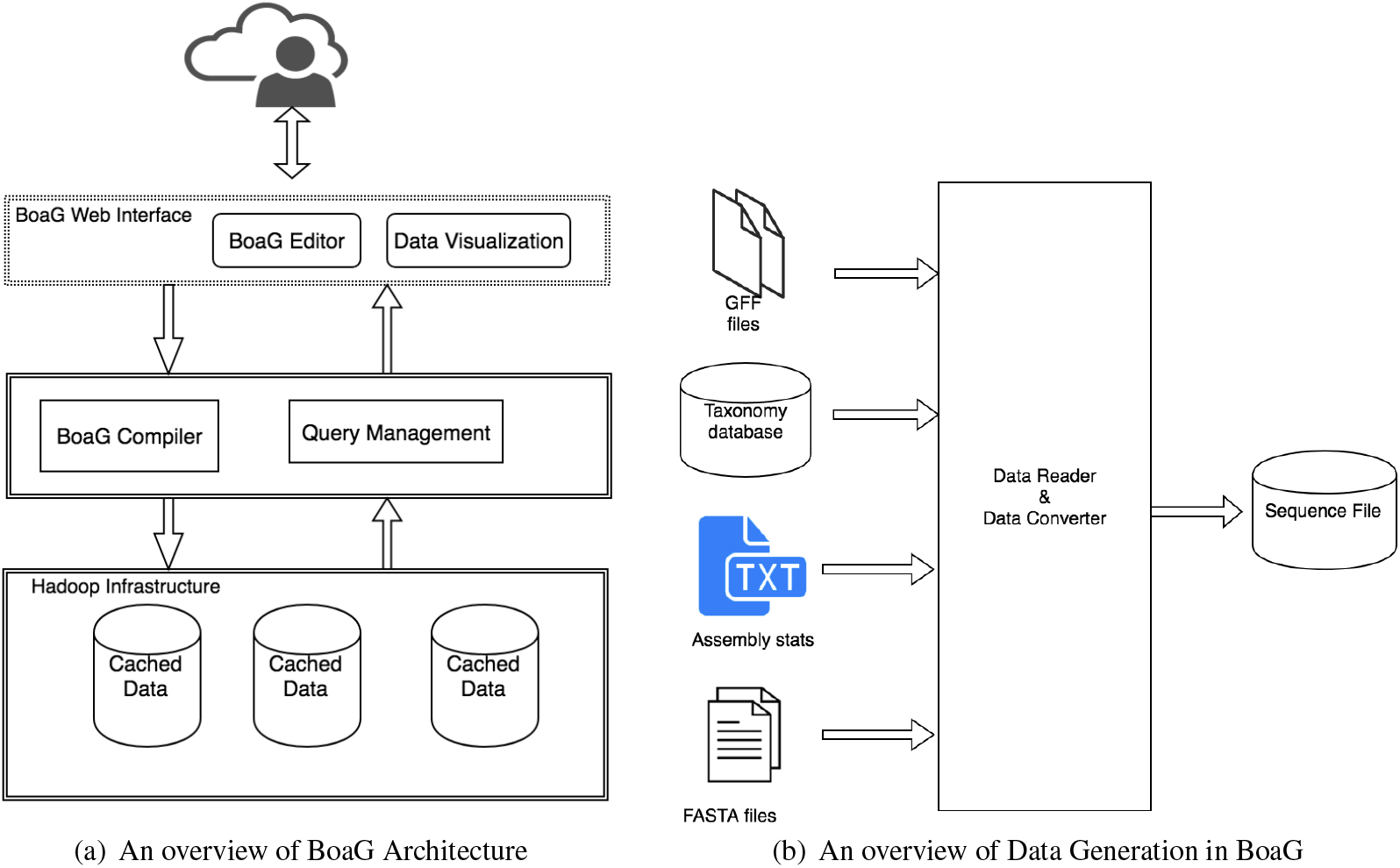
BoaG Architecture and Data Generation

#### Genomics-specific Language and data schema

To create the domain specific language for genomics in Boa, we created domain types, attributes and functions for this relatively small RefSeq dataset that includes the following raw file types: FASTA, GFF and associated metadata, as shown in Figure 2. Genome, Sequence, Feature, and Assembler are types in BoaG language and taxid, refseq, etc are attributes of Genome type. We created the data schema based on the Google protocol buffer which is an efficient data representation of genomic data that provides both storage efficiency and efficient computation in BoaG.

**Figure 2:**
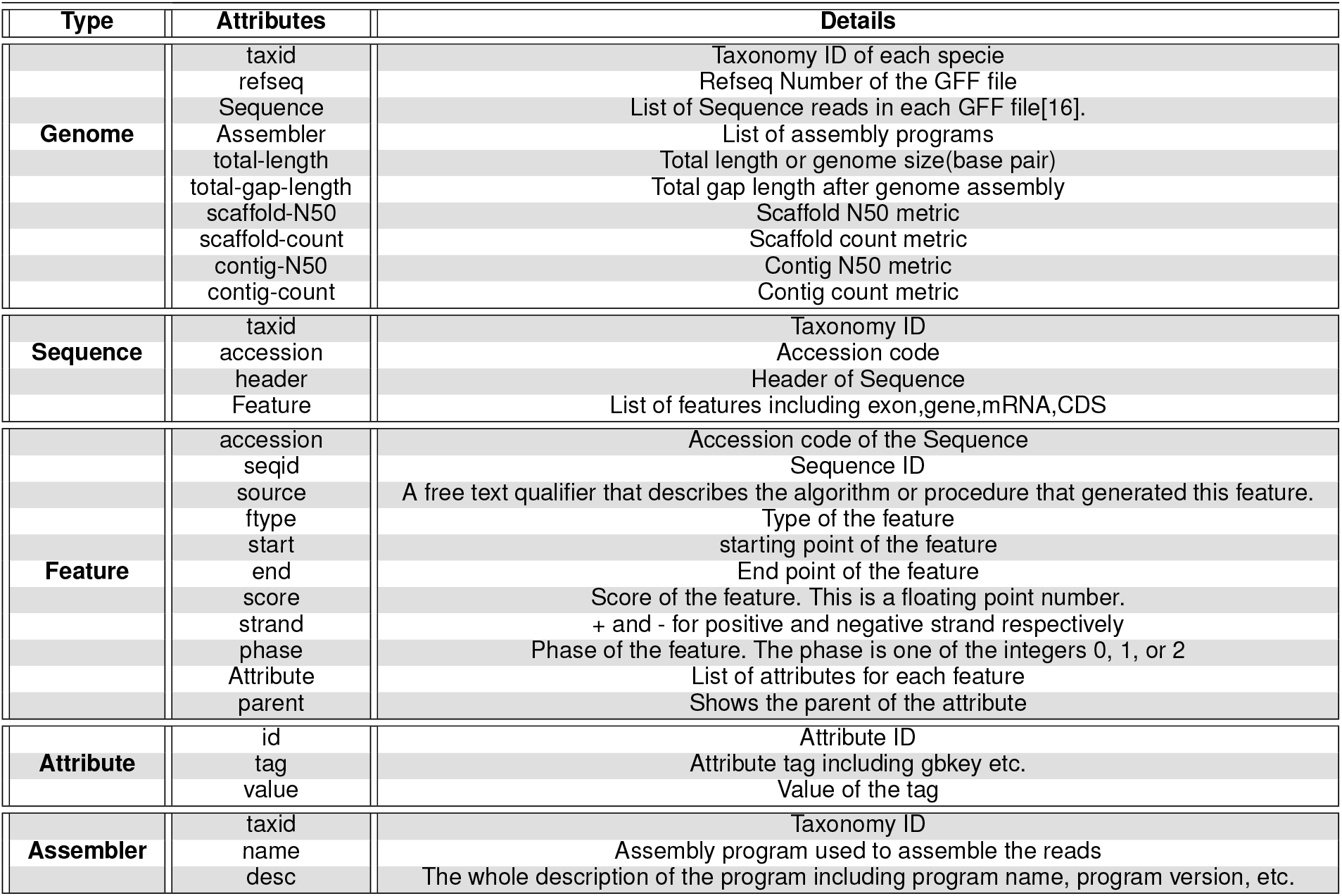
Domain types for Genomics data in BoaG

### BoaG database, Storage efficiency and Performance

The BoaG database is generated to fully utilize Hadoop. Raw data for each file type and metadata was parsed into a BoaG database (Hadoop sequence file) Figure 1(b). A compiler, file reader and converter was written in Java to generate this database and is provided on the GitHub repository (https://github.com/boalang/compiler). One benefit of the BoaG database is the significant reduction in required storage of the raw data. The NCBI RefSeq data presented here with a downloaded size of 167GB was reduced to 45.6GB (3.6 fold reduction) after writing to the BoaG database. A subset of the RefSeq data that contained only eukaryotic species showed an even more dramatic reduction from a raw data size of 58GB to a data size of 7.2 GB (8 fold reduction) once in the BoaG database format. The variability in size reduction is presumably due to variability in the number and size of files for eukaryotes vs bacteria. A second benefit of BoaG is its ability to take advantage of parallelization and distribution during computation, the greater the number of nodes the faster a BoaG program will complete. BoaG efficiency was tested on a Hadoop cluster with 40 nodes up to 400 map tasks. We ran different tasks on the preliminary RefSeq data on this cluster with varying levels of Hadoop mappers to show the dramatic speedup that results by adding additional Hadoop mappers to an analysis. As shown in Figure 3, four tasks were used to demonstrate the exponential decrease in required computation time with a corresponding increase in the number of Hadoop mappers. Specifically, Task1 was to determine the total number of genes for the 97,716 genomes contained in the RefSeq database, Task2 was to calculate the average summary statistics for every genome assembly, Task3 was to calculate the number of genes for every individual genome and Task4 was to determine which genome was the smallest and which genome was the largest in RefSeq (Rice yellow mottle virus satellite and the Aardvark, respectively)

**Figure 3:**
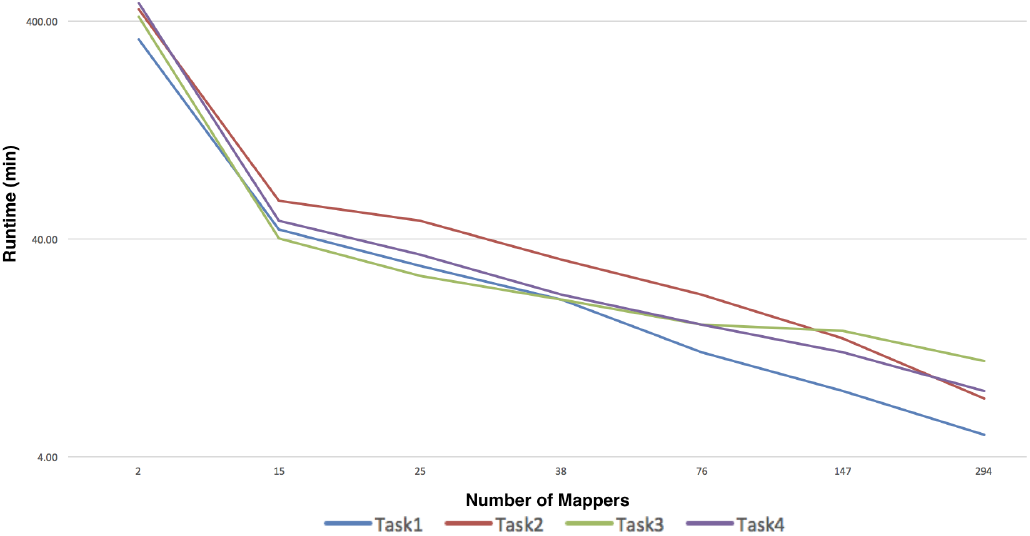
Scalability of BoaG programs

### Application of BoaG to the RefSeq database

A total of 97,716 annotation (GFF) and assembly (FASTA) files along with their associated metadata were downloaded from NCBI RefSeq [17] and written to a BoaG database. Metadata included genome assembly statistics (Genome size, scaffold count, scaffold N50, contig count, contig N50) and assembly program used to generate the assembly from which the genome annotation file was created. We were able to use BoaG to easily query the entire dataset and gain insight into the RefSeq genome assemblies and gene model annotations.

## Results and discussion

### Exploring RefSeq with BoaG

The downloaded RefSeq data consists primarily of bacterial genomes (89,195), followed by archaea (597) and then animal (363), fungal (237) and plant (84). Each genome has metadata related to the quality of its assembly (Genome size, scaffold count, scaffold N50, contig count, contig N50) and the assembly program used for the assembly. While it is straightforward to use the RefSeq website (https://www.ncbi.nlm.nih.gov/refseq/) to look up this information for your favorite species, it is cumbersome to look up this information for tens to hundreds or even thousands of related species. Similarly, while each of these genomes have an annotation file, querying and summarizing information contained in this annotation file from several related genomes such as average number of genes, average number of exons per gene and average gene size requires downloading and organizing the annotation files of interest prior to calculating gene model statistics. With BoaG it is relatively straightforward to answer these and many more questions of the RefSeq data we downloaded.

### Genome Assembly statistics varies across kingdoms

There were no surprises in the average size of genomes assembled in the five kingdoms: bacteria, archaea, fungi, plants and animals. As shown in Table 1, bacteria are around 3.6 Mb in size, archaea 2.6 Mb, fungi around 26 Mb, plants around 381 Mb and animals around 1170 Mb. As expected, in both plants and animals there were large standard deviation in the genome size indicating a large range of genome sizes in those kingdoms. Interestingly, we see that despite the popularity in recent years with long read technology and the ability to get closed genome assemblies for prokaryotes (bacteria and archaea) we still see a fairly high level of fragmentation in prokaryotes found in the RefSeq database with an average scaffold count of 53 scaffolds. It is also striking to see the difference in the fragmentation between plant and animal genomes: N50:1.5Mb vs N50:18.2Mb suggesting that animal genomes in RefSeq are on average better assembled. At the broad kingdom level, summary statistics has very little practical use to researchers. However, as we explore subclades we find some interesting trends. For example, different groups of closely related species tend to favor one particular assembly program over another. Is this due to groups of researchers that study different species have different preferences or is it that on their species they get a better assembly using one assembly program over another. Table 1 suggests it is the latter and was previously suggested by the Assemblathon paper [18] due to differences in genomic characteristics, repetitiveness, heterozygosity, ploidy, GC content, raw data, and so forth.

**Table 1:** Traditional kingdoms and average summary statistics for their genome assemblies

| Tax ID | Name | Species | Total length | Scaffold-count | ScaffoldN50 | ContigCount | ContigN50 |
| --- | --- | --- | --- | --- | --- | --- | --- |
| 2 | Bacteria | 89195 | 3.6M ± 1.6M | 53.5 ± 114.6 | 1.5M ± 1.5M | 119.3 ± 185 | 244.1k ± 561k |
| 4751 | Fungi | 236 | 26.2M ± 19.9M | 265.8 ± 361 | 2M ± 1.5M | 890.1 ± 1742 | 704.8k ± 886k |
| 2157 | Archaea | 597 | 2.6M ± 1M | 11.1 ± 11.9 | 1.6M ± 1M | 81.9 ± 155 | 525.8k ± 615k |
| 33090 | Viridiplantae | 84 | 381.8M ± 453.2M | 8143 ± 11k | 1.9M ± 1.1M | 24.4k ± 28.8k | 1.3M ± 3.3M |
| 33208 | Metazoas | 363 | 1.17B ± 1.15B | 25.4k ± 55.2k | 18.2M ± 25.2M | 79.2k ± 100k | 2.28M ± 9.4M |
| 71240 | eudicotyledons(dicots) | 57 | 636.7M ± 554.7M | 15.2k ± 13.5k | 2.5M ± 1.4M | 38.3k ± 34.2k | 1.64M ± 3.89M |
| 4447 | Liliopsida(monocotyledons) | 14 | 423.32M | 7512 | 1.31M | 14.77k | 3.86M |

For example, consider Bacteroidetes (TaxID 976) and Insecta (TaxID 50557). As it can be seen in Table 2, for insects the two most popular assembly programs are SoapDenovo and Allpaths while for Bacteroidetes the most popular assembly programs are Allpaths and Velvet. In both cases, Allpaths appears to do a better job at genome assembly with fewer scaffolds and a larger N50 value. However for bacteria, Newbler, the third most popular assembler may be slightly better in Bacteroidetes with comparable number of scaffolds and a considerably larger N50 value. As you can imagine, being able to mine RefSeq in this way would be a valuable tool to researchers to determining the best choice of assembly program to start with for a particular species and a means to compare their assembly quality with tens to hundreds of related species relatively quickly. In the coming years as more species are sequenced, this ability will be very important for the accurate assessment of the quality of an assembly.

**Table 2:** List of top five most used assembly programs for Insects and Bacteria

| Kingdom | Program Name | species | Total length | Scaffold-count | ScaffoldN50 | ContigCount | ContigN50 |
| --- | --- | --- | --- | --- | --- | --- | --- |
| Bacteria | Allpaths | 8846 | 4.40M ± 1.40M | 28.68 ± 32.14 | 1.46M ± 1.59M | 103.08 ± 126.80 | 366.81k ± 658.57k |
|  | Velvet | 4380 | 3.83M ± 1.56M | 92.72 ± 121.53 | 861.66k ± 1.52M | 130.95 ± 175.19 | 246.72k ± 537.16k |
|  | Newbler | 3517 | 4.24M ± 1.53M | 33.34 ± 57.91 | 2.25M ± 1.92M | 122.7 ± 142.3 | 295.88k ± 604.01k |
|  | CLC | 2274 | 3.93M ± 1.53M | 40.30 ± 64.58 | 2.23M ± 1.96M | 130.77 ± 173.09 | 235.47k ± 492.06k |
|  | Spades | 2013 | 4.04M ± 1.56M | 38.93 ± 60.09 | 2.17M ± 1.92M | 127.56 ± 168.95 | 246.85k ± 506.66k |
| Insects | Soapdenovo | 26 | 323M ± 183.22M | 21.12k ± 40.29k | 1.46M ± 1.81M | 43.74 ± 44.58 | 40.03k ± 36.13k |
|  | Allpaths | 19 | 353.69M ± 260.41M | 3821 ± 3778 | 2.41M ± 1.94M | 34.14k ± 35.37k | 40.95k ± 23.48k |
|  | Celera | 10 | 179.96M ± 30.36M | 6270 ± 2231 | 1.14M ± 961.18k | 9354 ± 5393 | 292.96k ± 257.23k |
|  | Newbler | 8 | 305.55M ± 123.14M | 22.79k ± 21.15k | 1.88M ± 1.11M | 39.52k ± 29.36k | 48.02k ± 30.83k |
|  | Atlas | 7 | 540.68M ± 361.56M | 4832 ± 4146 | 5.05M ± 4.39M | 53.7k ± 47.04k | 67.38k ± 71.52k |

It is important to mention here that an analysis in BoaG requires fewer lines of codes. For example, as shown in Figure 4, the top three most used assembly program required only 5 lines in the BoaG language to answer this question whereas a similar analysis using python required 38 lines of code (supplemental file Figure 5). This advantage inherent to domain specific languages will speed up a researcher’s ability to query large datasets.

**Figure 4:**
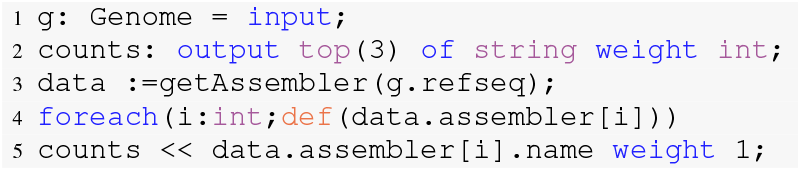
What are the top three most used assembler programs?

### Genome Annotation statistics and average exons per gene

Using BoaG we calculated average summary statistics for the gene model annotations of the genomes in each of the specie’s clades in RefSeq. The summary statistics we chose, as shown in Table 5, included gene number, exon number and number of exons per gene. As we would expect due to the smaller genome size, we find fewer genes in prokaryotes (bacteria and archaea) with 3,982 and 2,864 respectively than Eukaryotes (plant, animal and fungi) with 34,460, 22,700 and 9,472 respectively.

As is common knowledge, number of exons in prokaryotes is equivalent to the number of genes since prokaryotes do not have introns and the number of exons per gene in our analysis is equal to 1. As it can be seen in Table 3, the average number of exons per gene in fungi (3.66 ± 1.59) is lower than that in plants (6.54 ± 1.78) and animals (7.54 ± 1.5). With BoaG and the RefSeq database it is easy to identify the top 10 genomes with the highest and lowest average exons per gene in Eukaryotes. The genomes with the highest average number of exons per gene are primarily in the animal kingdom with nine of the ten top ten genomes belong to this kingdom (Table 4). This is inline with our results that indicate Metazoans (animals) have higher average exons per gene than plants and fungi. The one outlier is a fungus (Serpula lacrymans). This outlier may be an artifact of poor annotation or may indicate some evolutionary advantage for higher average exon number per gene. Further investigation is warranted but is beyond the scope of this work. Similarly, as it can be seen in Table 5, the top 10 Eukaryotic genomes with the lowest average exons per gene were all fungi which have similar levels to that of prokaryotes with close to 1 exon per gene on average. Why do fungi have a lower number of exons per gene than other prokaryotes and which fungi have more and what evolutionary advantage does this provide are all questions that are beyond the scope of this work but demonstrate the inherent worth of being able to explore data repositories using a data science infrastructure like BoaG.

**Table 3:**
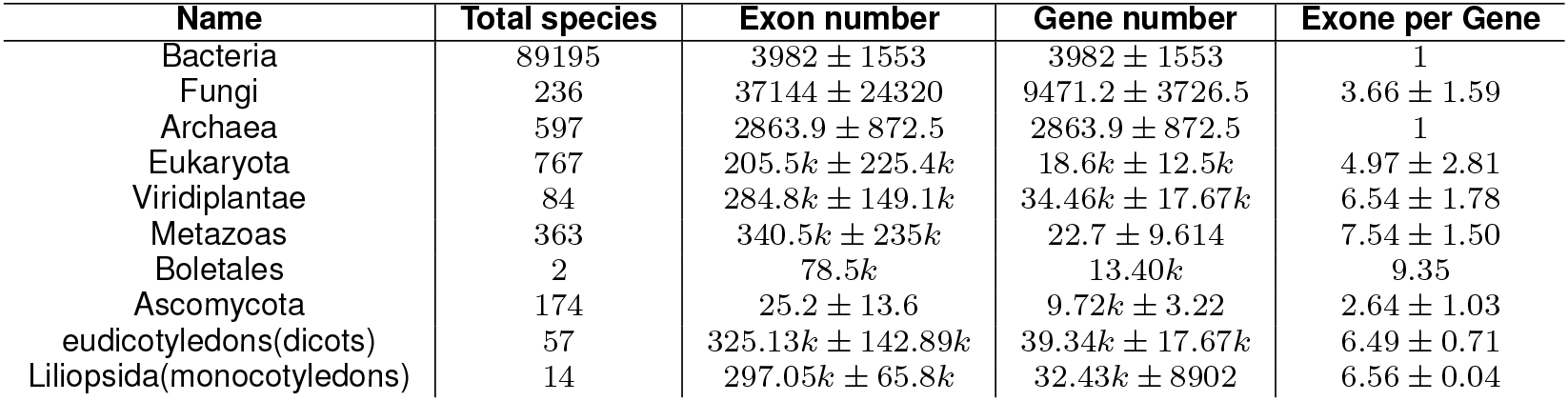
Exon Statistics

| Name | Total species | Exon number | Gene number | Exone per Gene |
| --- | --- | --- | --- | --- |
| Bacteria | 89195 | 3982 ± 1553 | 3982 ± 1553 | 1 |
| Fungi | 236 | 37144 ± 24320 | 9471.2 ± 3726.5 | 3.66 ± 1.59 |
| Archaea | 597 | 2863.9 ± 872.5 | 2863.9 ± 872.5 | 1 |
| Eukaryota | 767 | 205.5k ± 225.4k | 18.6k ± 12.5k | 4.97 ± 2.81 |
| Viridiplantae | 84 | 284.8k ± 149.1k | 34.46k ± 17.67k | 6.54 ± 1.78 |
| Metazoas | 363 | 340.5k ± 235k | 22.7 ± 9.614 | 7.54 ± 1.50 |
| Boletales | 2 | 78.5k | 13.40k | 9.35 |
| Ascomycota | 174 | 25.2 ± 13.6 | 9.72k ± 3.22 | 2.64 ± 1.03 |
| eudicotyledons(dicots) | 57 | 325.13k ± 142.89k | 39.34k ± 17.67k | 6.49 ± 0.71 |
| Liliopsida(monocotyledons) | 14 | 297.05k ± 65.8k | 32.43k ± 8902 | 6.56 ± 0.04 |

**Table 4:** Eukaryotic genomes with the top 10 highest average number of exons per gene

| Scientific Name | Tax id | Exon number | Gene number | Exon per gene |
| --- | --- | --- | --- | --- |
| Branchiostoma belcheri | 7741 | 416715 | 26845 | 11.94 |
| Crassostrea gigas | 29159 | 523578 | 33802 | 11.48 |
| Actinobacillus hominis | 7719 | 216702 | 15301 | 11.15 |
| Cimex lectularius | 79782 | 261455 | 14888 | 10.49 |
| Lingula anatina | 7574 | 448972 | 32398 | 10.42 |
| Loa loa | 7209 | 105705 | 15040 | 10.22 |
| Serpula lacrymans var. lacrymans S7.9 | 578457 | 72891 | 12923 | 10.19 |
| Nematostella vectensis | 45351 | 133623 | 27173 | 10.14 |
| Halyomorpha halys | 286706 | 252938 | 16620 | 9.91 |
| Polistes dominula | 743375 | 210267 | 11313 | 9.88 |

**Table 5:**
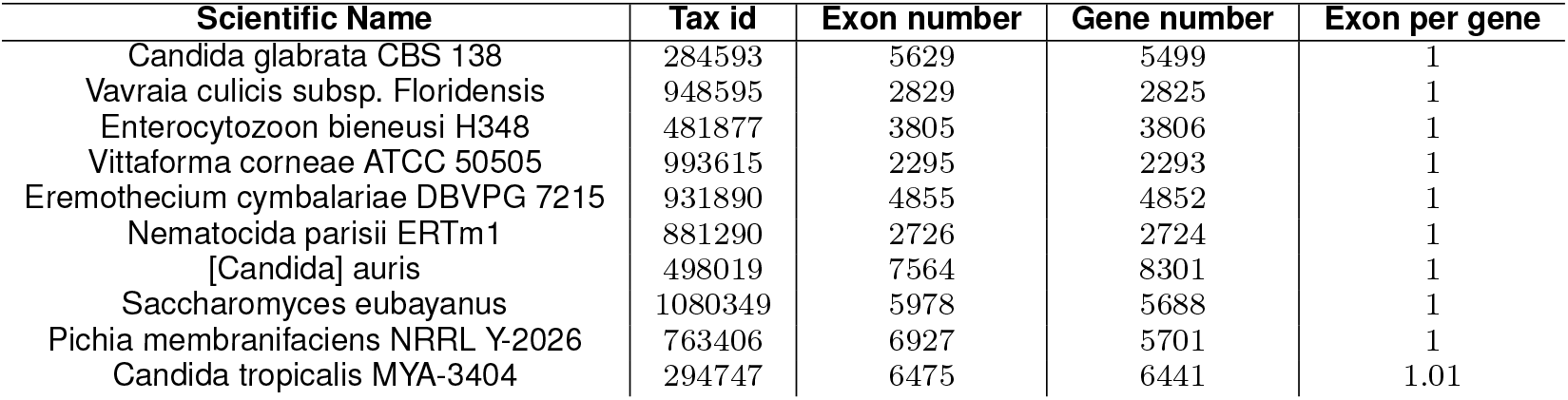
Eukaryotic genomes with the top 10 lowest average number of exons per gene

### Future work

Being able to efficiently query FASTA and GFF files is only the smallest part of Boa’s capabilities. In the future, the BoaG infrastructure will be extended to support more data files by providing data reader and data converter. For each data file, new DSL types will be defined and the BoaG compiler also will be updated to read new DSL types. Furthermore, functions will be added in the BoaG domain language and infrastructure. Some functions that would be directly relevant to the our RefSeq dataset include direct calculation of assembly statistic and BUSCO scores. Since potential issues with input data format are taken care of during the parsing and writing of database, the function written in BoaG will work on every raw data file in the dataset. Furthermore, since the BoaG infrastructure operates on top of a Hadoop cluster, all functions generated will also be extremely scalable. Using the RefSeq data as an example, it is not difficult to imagine building an interface to the BoaG infrastructure in the form of a website that would allow researchers to query genome assembly and annotation statistics data to determine the relative quality of their newly assembled genome with closely related assembled genomes for any clade or subset of genomes in RefSeq. In addition, the website could work as an interface for users to execute BoaG scripts directly on a Hadoop cluster.

## Conclusion

In this work, we presented BoaG which is a domain specific language and shared data science infrastructure that takes advantage of Hadoop distribution for large-scale computations. Its infrastructure opens up possibilities to explore data in ways previously not possible without deep expertise in data acquisition, data storage, data retrieval, data mining and parallelization. BoaG was effectively used to explore the RefSeq database of genome assemblies and annotations to identify interesting features of gene annotation specific to individual clades. While the simple query examples discussed here could also have been performed using traditional databases, it provides a proof of concept behind the BoaG infrastructure and its ability to scale to much larger datasets.

## Additional Files

Additional file 1 — BoaG comparison with Python

Figure 5 Compare Line of Code(LOC) and performance to answer query “ What are the top three most used assembly program?” run on Refseq Data. The BoaG run time and numbers of lines of codes with the corresponding python run time and codes represent the BoaG significant improvements.

**Figure 5:**
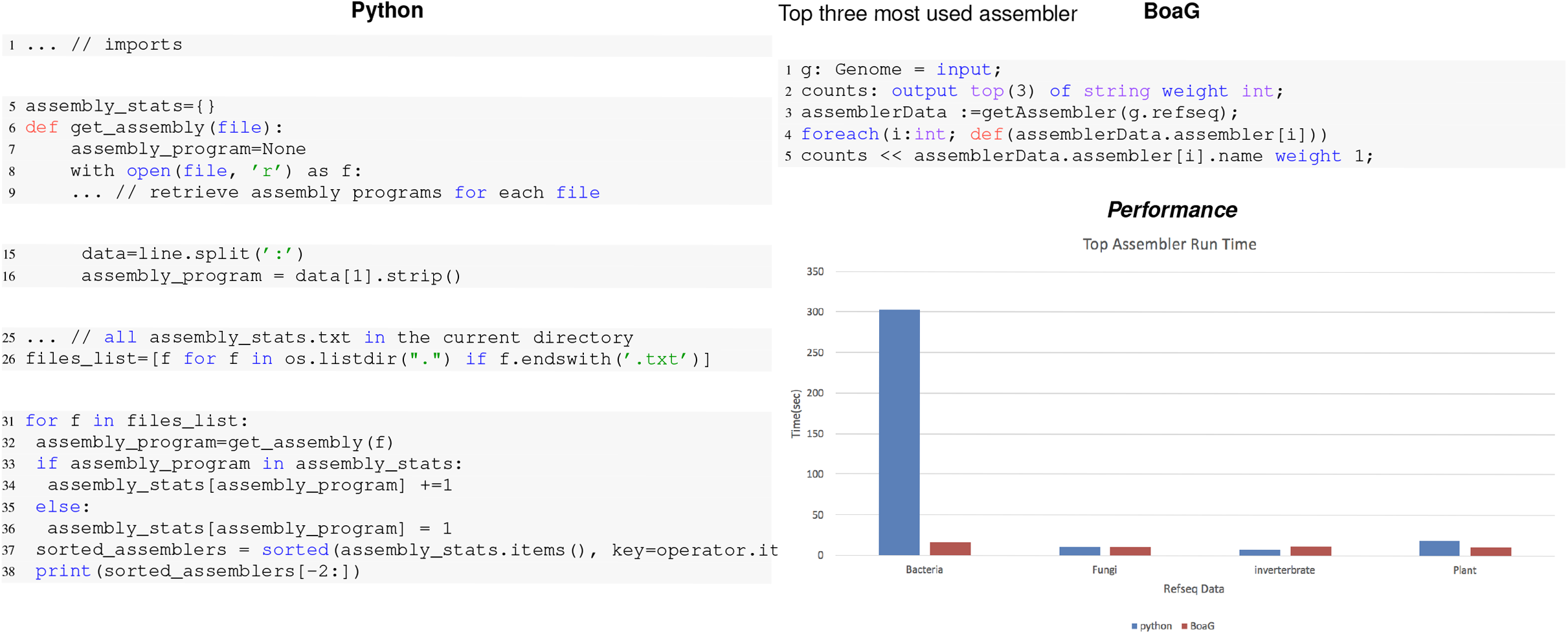
Comparison of Line of Code(LOC) and performance to answer query “ What are the top three most used assembly program?” run on Refseq Data. On the left side, the equivalent BoaG code needs 38 lines of code in Python whereas the BoaG script needs only 5 lines of code.

## Abbreviations

BoaG: Boa for Genomics; SDSI: Shared Data Science Infrastructure

## Ethics approval and consent to participate

Not applicable.

## Consent for publication

Not applicable.

## Availability of data and material

Boa compiler’s source code is provided on the GitHub repository (https://github.com/boalang/compiler)

## Author’s contributions

Severin and Muppirala conceived of the application of Boa to the RefSeq database and contributed to exploring the dataset from a biological perspective. Hamid Bagheri wrote the codes, implemented the genomics specific types and customized compiler. He ran analysis and prepared figures. Muppirala provided a first outline of the paper that Bagheri later fleshed out before additional rounds of major editing by all authors. Hridesh Rajan invented the notion of Shared Data Science Infrastructure (SDSI) [19], and the idea of SDSI for genomics data. He also contributed to the design of the BoaG domain-specific language for computing over data, and domain-specific types for representing RefSeq data. All the authors read and approved the final manuscript.

## Acknowledgments

We would like to thank the VPR office at Iowa State University to provide seed funding to explore the application of Boa to the biological domain. This material is based upon work supported by the National Science Foundation under Grant CCF-15-18897 and CNS-15-13263. Any opinions, findings, and conclusions or recommendations expressed in this material are those of the authors and do not necessarily reflect the views of the National Science Foundation

